# Conditional genome-wide associations reveal novel genes

**DOI:** 10.64898/2026.04.08.717258

**Authors:** Emily S. Bellis, Marta Robertson, William W. Booker, Cynthia Rudin, Mariano F. Alvarez

**Author notes:** Corresponding author Correspondence to: Mariano Alvarez. Contributions M.F.A. developed the algorithms, with input from C.R. M.F.A., E.S.B and W.W.B. conducted computational experiments. E.S.B. and M.R. designed validation experiments, and M.R. analyzed experiment results. E.S.B. drafted the manuscript with contributions from all authors. All authors contributed to review and revision.

## Abstract

We introduce two novel approaches for gene discovery based on conditional genome-wide associations. Experimental validation of gene targets identified by our top-performing approach uncovers three genes with a previously unknown role in controlling flowering time in *Arabidopsis*, one of the most well-studied traits in the most well-studied plant genome. This work demonstrates the power of knockoff-based frameworks to uniquely identify novel genes underlying complex traits, a core task across applications in agriculture and human health.

## Main

Understanding the genetic basis of complex traits has remained a foremost goal in the biological sciences over the past few decades. Yet, large portions of gene space remain fully uncharacterized across the tree of life. Across the human genome, for example, more than a fifth of protein families still have a completely unknown function^1^. Genome-wide association studies (GWAS) have served as a keystone methodology for linking genes to variation in complex traits, with a standard GWAS capable of producing hundreds or thousands of testable hypotheses of gene function. However, many GWAS hits end up being false discoveries, and overall, tend to explain only a fraction of genetic variance in traits. This ‘missing heritability’ problem is thought to stem in large part from a multitude of small effect variants across the genome, which together contribute a major portion of trait variance but are very difficult to detect using traditional GWAS methods^2,3^.

Recent developments in the field of explainable machine learning have shown improvements over traditional inference methods for gene discovery from genome-wide data. In particular, the knockoff filter^4^, which strictly controls the false discovery rate (FDR) based on comparisons to synthetically generated variables, is agnostic of the underlying model architecture linking the explanatory variables with the response and has been useful to address confounding due to genetic linkage and population structure^5,6^. While knockoffs, which can be based on fundamentally more flexible models than widely used generalized linear mixed model (GLMM)-based approaches, are now being applied more broadly beyond humans, most authors to-date validate these approaches based on the ability to recover known genes discovered with traditional methods. Such a strategy provides little support for the idea that knockoffs and related methods should be well-suited to reveal *novel* genome associations.

Here, we present two novel approaches for genomic feature selection and gene discovery. Our strategy, based on principles for conditional model reliance (CMR)^7^, is conceptually similar to knockoffs, in that the predictive power of a focal variable is assessed by replacing it with a synthetic variable. Unlike existing knockoff frameworks, the synthetic variable is generated such that it carries only information about the dependent variable that can be gleaned from other covariates. In other words, by removing a SNP and then constructing a new one using information from a generative model created without information from the removed SNP, we can generate a synthetic SNP that contains all information from the model except the information unique to that position. Rather than testing the resilience of the variable to perturbation, our CMR-based algorithms instead allow us to directly test the importance of the conditional value. That is, we assess how valuable the information unique to that variable is.

Our gene discovery through information-less perturbation (GDIP) approach includes the following main steps: 1) generate one or more knockoffs for each variant, 2) calculate a feature importance score for each original feature and their associated knockoff(s), and 3) calculate a test statistic based on the divergence between the importance scores of the original feature and the knockoff(s). In contrast to existing knockoffs-based frameworks, we generate knockoffs for each variant based on conditional model reliance (CMR)^7^. Considering a sample population of *n* individuals genotyped at *p* genetic markers:

For *j*= 1, …, *p*:

Let ***X***_−*j*_ denote the feature matrix ***X*** ∈ ℝ ^*n*×*p*^ excluding column *j*

Construct CMR-based knockoff 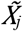 using only ***X***_−*j*_ such that:

a. 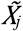 preserves correlation structure with ***X***_−*j*_
b. 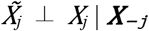

Following He et al.^20^ we also developed a version of GDIP based on summary statistics, referred to throughout the manuscript as GDIP-gk. Just as for GDIP, GDIP-gk removes information about the feature of interest from the knockoff generation procedure as follows:

For *j*= 1, …, *p*:

Let ***z***_−*j*_ denote a *p-*dimensional vector of *Z*-scores excluding element *j*

Sample knockoff statistic 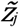 using only ***z***_−*j*_ such that:

a. 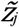 preserves dependency structure with ***z***_−*j*_
b. 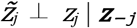

Use *z*_*j*_ and 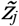 as importance scores for original and knockoff features, respectively.

We benchmarked our CMR-based knockoff approaches (GDIP and GDIP-gk) on simulated data against existing GWA methods based on group knockoffs (SNPknock)^8^ and GLMMs (GEMMA)^9^ (Fig. 1). For simulated ‘easy’ traits (low polygenicity and medium effect sizes), we found that all knockoff-based methods including our CMR-based approaches dramatically outperformed linear models. Both of our CMR-based approaches were characterized by significantly higher recall compared to both SNPknock and GEMMA. On simulated ‘challenging’ traits (high polygenicity with weak effect sizes), performance of all methods worsened substantially. However, both of our CMR-based approaches maintained their advantage of higher recall compared to SNPknock and GEMMA, with GDIP-gk demonstrating less variability but also higher precision than GDIP. Overall, GDIP-gk was the top-performing method according to the F1 score, with a median improvement from 1.6x - 2.4x over SNPknock depending on the task (Fig. 1).

**Figure 1.**
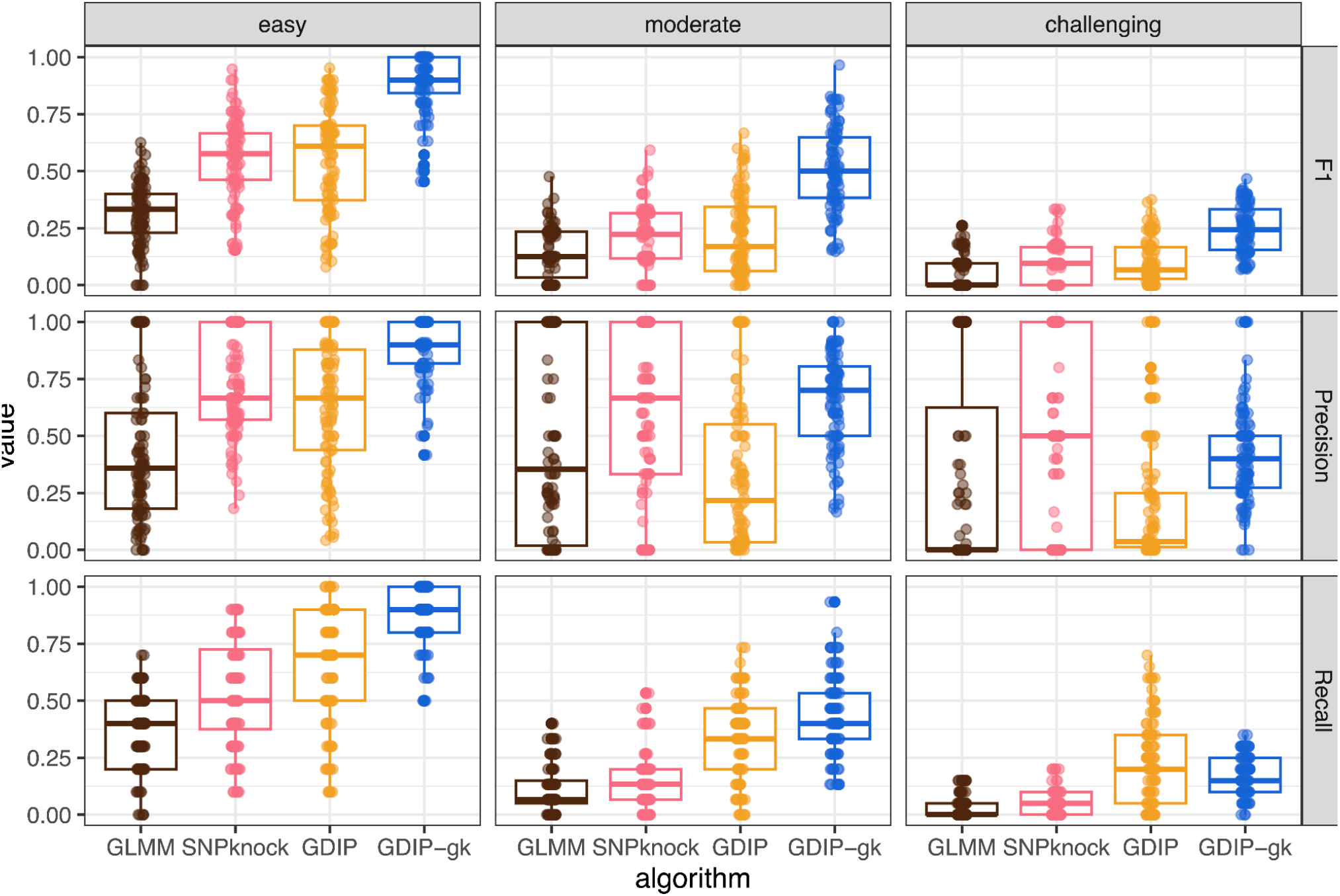
Evaluation of gene discovery methods on simulated traits. F1 score, precision, and recall metrics for 100 simulated traits per task (‘easy’, ‘moderate’, or ‘challenging’). Metrics are reported for a generalized linear mixed model (GLMM), an existing knock-offs-based method (SNPknock), and two versions of our GDIP algorithm.

To further validate our top performing method, we carried out a GWAS for flowering time using an existing dataset from the 1001 Arabidopsis Genomes Project and our top-performing algorithm (GDIP-gk). A similar, small proportion of previously known flowering time genes was recovered by GDIP-gk vs. GLMM (Fig. 2H). Among GDIP-gk hits, we identified 69 sites significant at an FDR of 0.1, which tagged 153 genes within 5kb. Of these 153 genes, only four (2.6%) were present among the reference set of 306 known flowering genes including *FLOWERING LOCUS T* (*FT*; AT1G65480), *COLD, CIRCADIAN RHYTHM, AND RNA BINDING 2* (*CCR2*; AT2G21660), *ZEITLUPE* (*ZTL;* AT5G57360), and *VERNALIZATION INSENSITIVE 3* (VIN3; AT5G57380) (Fig. 2a). By comparison, 589 GLMM hits tagged 282 genes, of which seven (2.5%) were present in the reference set. In addition to *FT, ZTL*, and *VIN3*, which were also found by GDIP-gk, the four additional true positive genes implicated by GLMM included *CRYPTOCHROME-INTERACTING BASIC-LOOP-HELIX 5* (*CIB5*; AT1G26260), *TWIN SISTER OF FT* (*TSF*; AT4G20370), *FLOWERING LOCUS C* (*FLC;* AT5G10140), and *CONSTANS-LIKE 5* (*COL5*; AT5G57660). Two of these genes (*FLC* and COL5) were also tagged by GDIP-gk but by a stronger association signal beyond our 5 kb cut-off. Specifically, *FLC* was tagged by a significant GDIP-gk hit 8.9 kb away, and this site had the strongest GLMM association to flowering time in a 40 kb region of extended LD around *FLC* (Fig. 2A). The closest GDIP-gk hit to *COL5* was 66.6 kb upstream but also demonstrated stronger associations to flowering time than other GLMM hits in the region. Although the proportion of genes from the reference set was not statistically different between methods, hits identified by GDIP-gk were >4x more likely to uniquely tag a gene model (0.45 hits/gene) compared to GLMM (2.1 hits/gene), highlighting the large number of redundant hypotheses associated with the latter method.

**Figure 2.**
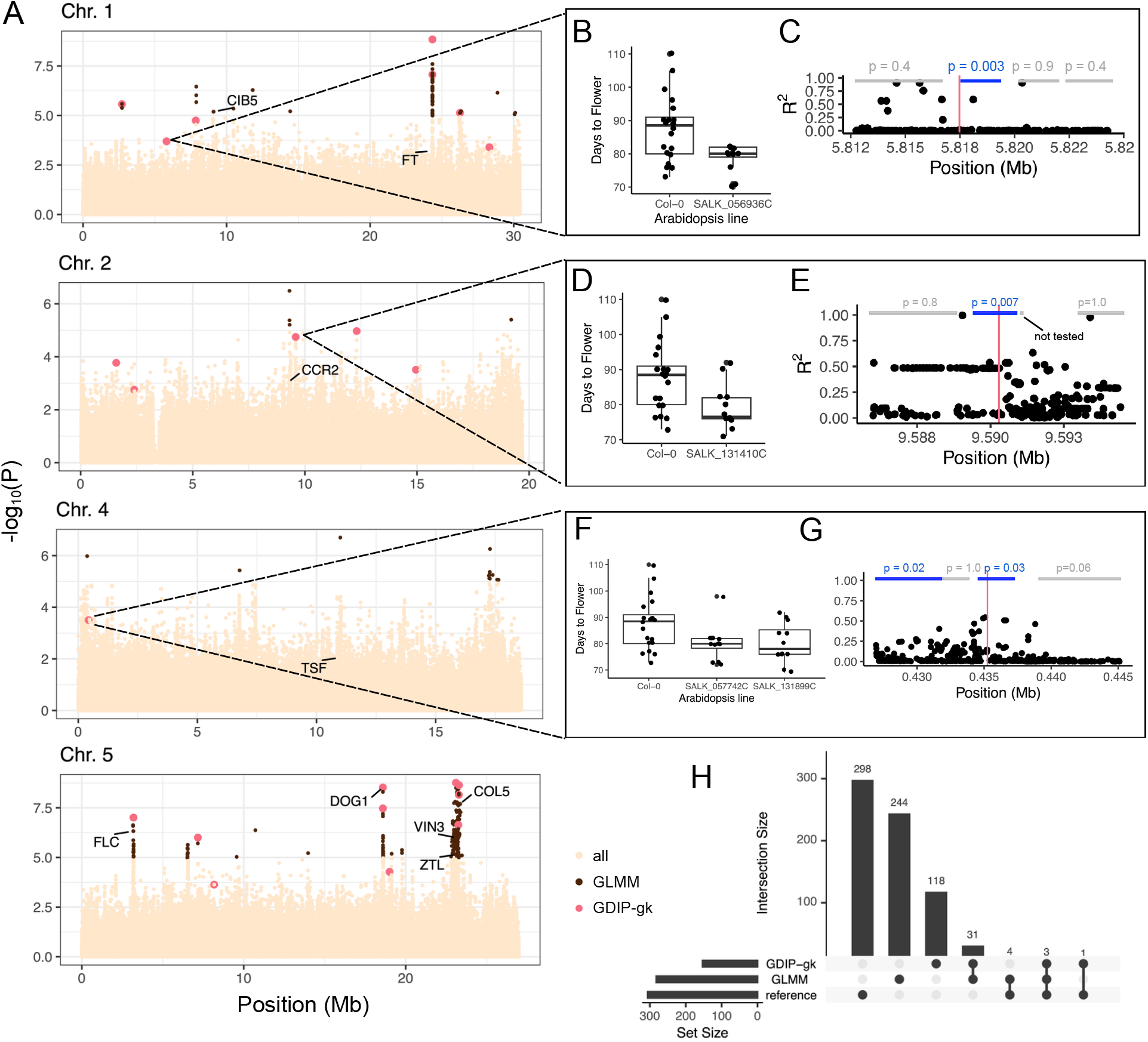
Experimental validation of GDIP-gk hits. (A) Arabidopsis genome-wide associations for flowering time at 10°C. SNPs are plotted according to associations determined from the GLMM, with significant GDIP-gk hits at an FDR threshold of 0.1 colored in pink. Positions of known flowering time genes are shown. (B, D, F) Days to flowering for Arabidopsis T-DNA mutant lines with significantly earlier flowering than Col-0. (C, E, G) Linkage disequilibrium for SNPs within the boundaries of tested gene models. The location of the focal GDIP-gk hit is shown by the vertical pink line. Boundaries of experimentally evaluated gene models are shown by blue rectangles, with the *p-*value for the comparison of the corresponding T-DNA insertional mutant line vs. Col-0. (H) Overlap among sets of genes tagged by GDIP-gk, a GLMM, or present in a reference set of 306 known flowering time genes. Most genes were unique to a set.

By reducing the number of redundant hypotheses tested in a GWAS, knockoff-based approaches should be well suited to identify novel hits that would fail to meet a significance threshold with existing FDR control procedures. We tested this idea through experimental follow-up of 11 GDIP-gk hits for *Arabidopsis* flowering time that did not meet the significance threshold for GLMM. Across 28 gene models within 5-kb of one of these 11 GDIP-gk hits, insertional mutant lines for four genes showed significantly earlier flowering compared to wild-type (Col-0). On Chromosome 1, we tested 4 T-DNA mutant lines for gene models within 5-kb of a hit at position 5,817,473 (MAF: 2%; Fig. 2B-C) and found that disruption of AT1G17010 (described as a 2-oxoglutarate and Fe(II)-dependent oxygenase superfamily protein) resulted in 9.5 day earlier flowering compared to Col-0 (*p =* 0.003). AT1G17010 exhibits high gene expression specifically in seeds^10^, suggesting a potential contribution to coordination of seasonal life-history strategies at low temperature^11^. On Chromosome 2, we tested 3 T-DNA mutant lines for genes within 5-kb of position 9,590,291 (MAF: 48%; Fig. 2D-E) and found that disruption of *NICOTINAMIDASE 1* (*NIC-1*; AT2G22570) resulted in early flowering of 9.0 days compared to Col-0 (*P =* 0.007). *NIC-1* may contribute to flowering time variation by controlling response to abscisic acid^12^. On Chromosome 4, we tested T-DNA mutant lines for 4 genes near a GDIP-gk hit at position 435,252 (MAF: 17%; Fig. 2F-G). We found that disruption of *JMJC DOMAIN-CONTAINING PROTEIN 27* (*JMJ27;* AT4G00990), with a previously reported role in flowering time^13^, resulted in early flowering of 8.1 days compared to Col-0 in our study (*P =* 0.025). Unexpectedly, a second T-DNA line with a disruption mutation in AT4G01010, which contained the GDIP-gk hit, also showed a significant 7.9 day decrease in flowering (*P =* 0.030). AT4G01010 encodes *CYCLIC NUCLEOTIDE-GATED CHANNEL 13* (*CNGC13*), which may contribute to flowering time regulation by modulating Ca^2+^ oscillations^14^.

Together, our experimental validation results support a key role in flowering time for 3 of 11 hits (27%) identified using our GDIP-gk and knockoff-based approach, but unidentifiable from a GLMM-based GWAS. A great deal of the ‘missing heritability’ in complex traits may be hiding in plain sight, but only discoverable with improved methods for gene discovery. Our work suggests that much remains to be learned about the genotype-phenotype map from the existing treasure trove of GWA datasets.

## Methods

### Simulated phenotypes

All simulations were conducted using SNP data derived from *A. thaliana*^15^ and supplied as part of the *Naturalgwas* package^16^. *Naturalgwas* generates synthetic phenotypes for existing genomic data, drawing from principal components to model population structure. Each synthetic phenotype is associated to randomly chosen causal loci, allowing for a fair comparison of power and false discovery between methods. We simulated one hundred independent phenotypes for the model plant *Arabidopsis thaliana* for each of three types of traits: 1) traits with an ‘easier’ genetic architecture for GWAS, 2) traits with a moderately challenging architecture, and 3) traits with a challenging genetic architecture. For our phenotype simulations representing an ‘easy’ GWAS task, 10 causal SNPs were drawn from unlinked loci, as described previously^16^, with effect sizes of each locus set to 100, representing a moderate effect size relative to the square root of the eigenvalues of the SNP matrix. For our phenotype simulations representing a ‘challenging’ GWAS task, we considered 20 causal SNPs and an effect size of 10. Finally, we also included a ‘moderate’ GWAS task with 15 causal SNPs and an effect size of 50. Each combination of parameters was represented in 100 separate simulations. Parameter values were chosen to span a full range of effect sizes and polygenicity as described previously^16^.

For each trait, we compared results from our GDIP and GDIP-gk algorithms against a GLMM-based model and an existing knockoffs-based approach. For the GLMM, we used GEMMA with a centered kinship matrix^9^, with FDR control using the Benjamini & Hochberg method^17^, implemented with the p.adjust function in R. For the existing knockoffs-based approach, we used SNPknock^8^ with group knockoffs constructed from unphased genotypes.

### Arabidopsis 1001 Genomes

We benchmarked GDIP-gk on real data using a genomic dataset of 5,775,084 single nucleotide polymorphisms (SNPs) and flowering time at 10°C from the 1001 Arabidopsis Genomes Project^18^. This dataset includes 1,003 natural inbred lines of Arabidopsis collected across its native range in Eurasia and North Africa.

### Experimental validation

From the all 69 regions identified by GDIP-gk significant at an FDR threshold of 0.1 (Supplemental Table S1), we excluded 13 also identified as significant in the GLMM-based analysis. We then manually selected 11 GDIP-gk regions for follow-up experimental validation (Supplemental Table S3), chosen to be reasonably evenly distributed across the genome. Each of the 11 regions was associated with at least one and up to four gene models that were within 5 kb either up or downstream of the GDIP-gk hit, totalling 32 genes. We obtained Arabidopsis T-DNA insertion mutant lines for 30 genes that were represented in the SALK collection and available from the Arabidopsis Biological Resource Center. After removing two lines with low germination (SALK_133903C and SALK_040805C), this left 28 total genes for evaluation.

Seeds were planted in Sungro Promix soil and stratified in the dark at 4C for 7 days. Plants were transferred to Percival LED-41L2 growth chambers and grown at constant 10°C and 16-hour daylength. Light intensity was 90 µmol·m^−2^·s^−1^. After germination, seedlings were thinned to 1 plant per well. Plants were grown in randomized complete blocks, with 1 replicate per block and 12 replicates per genotype total. The date of first flowering was recorded for each plant.

Time to flowering was analyzed with a linear mixed model (lme4 v 1.1-37, R version 4.4.2), with accession as a fixed effect and block as a random effect. Post-hoc comparisons were analyzed with estimated marginal means (R package emmeans v 2.0.0). A time-to-event analysis was also performed (R package survival v 3.7-0) with log-rank comparisons of each accession. The results of the two models were consistent, and the results of the linear mixed model are presented in text. Results of the survival analysis are presented in Supplementary Table 4.

## Data Availability

Phenotype data generated as part of this article for validation of the Salk T-DNA mutant lines are included in this published article and its supplementary information files.

## Code Availability

Code to demonstrate algorithm performance on a toy dataset is available on request to the corresponding author.

## Acknowledgements

We thank Michael Schwartz, Justin Sech, Emily Abernathy, Fared Farag, and John Joseph for their contributions to early experimental work and to development of computational infrastructure that supported the simulation benchmark experiments. We thank Shaoxing Huang, Tori Freeman, Leigh Schwinden, Emily Thayer, and Marisa Crisp for contributions to data collection. We thank the Salk Institute Genomic Analysis Laboratory for providing the sequence-indexed Arabidopsis T-DNA insertion mutants and Col-0.

## Supplementary Information

Supplemental Table S1: All significant GDIP-gk hits

Supplemental Table S2: GDIP-gk-tagged gene models

Supplemental Table S3: Experimental validation results

Supplemental Table S4: Survival Analysis Results

## References

1. Rocha, J. J. et al. Functional unknomics: Systematic screening of conserved genes of unknown function. PLoS Biol 21, e3002222 (2023).

2. Boyle, E. A., Li, Y. I. & Pritchard, J. K. An Expanded View of Complex Traits: From Polygenic to Omnigenic. Cell 169, 1177–1186 (2017).

3. Yang, J. et al. Common SNPs explain a large proportion of the heritability for human height. Nat Genet 42, 565–569 (2010).

4. Barber, R. F. & Candès, E. J. Controlling the false discovery rate via knockoffs. Ann. Statist. 43, (2015).

5. Sesia, M., Katsevich, E., Bates, S., Candès, E. & Sabatti, C. Multi-resolution localization of causal variants across the genome. Nat Commun 11, 1093 (2020).

6. Sesia, M., Bates, S., Candès, E., Marchini, J. & Sabatti, C. False discovery rate control in genome-wide association studies with population structure. Proc. Natl. Acad. Sci. U.S.A. 118, e2105841118 (2021).

7. Fisher, A., Rudin, C. & Dominici, F. All Models are Wrong, but Many are Useful: Learning a Variable’s Importance by Studying an Entire Class of Prediction Models Simultaneously. J Mach Learn Res 20, 177 (2019).

8. Sesia, M., Sabatti, C. & Candès, E. J. Gene hunting with hidden Markov model knockoffs. Biometrika 106, 1–18 (2019).

9. Zhou, X. & Stephens, M. Genome-wide efficient mixed-model analysis for association studies. Nat Genet 44, 821–824 (2012).

10. Klepikova, A. V., Kasianov, A. S., Gerasimov, E. S., Logacheva, M. D. & Penin, A. A. A high resolution map of the Arabidopsis thaliana developmental transcriptome based on RNA-seq profiling. The Plant Journal 88, 1058–1070 (2016).

11. Martínez-Berdeja, A. et al. Functional variants of DOG1 control seed chilling responses and variation in seasonal life-history strategies in Arabidopsis thaliana. Proc. Natl. Acad. Sci. U.S.A. 117, 2526–2534 (2020).

12. Wang, G. & Pichersky, E. Nicotinamidase participates in the salvage pathway of NAD biosynthesis in Arabidopsis. The Plant Journal 49, 1020–1029 (2007).

13. Dutta, A., Choudhary, P., Caruana, J. & Raina, R. JMJ 27, an Arabidopsis H3K9 histone demethylase, modulates defense against Pseudomonas syringae and flowering time. The Plant Journal 91, 1015–1028 (2017).

14. Dietrich, P. & Moeder, W. Plant Cyclic Nucleotide-Gated Channels: New Insights on Their Functions and Regulation. Plant Physiol. 184, 27–38 (2020).

15. Atwell, S. et al. Genome-wide association study of 107 phenotypes in Arabidopsis thaliana inbred lines. Nature 465, 627–631 (2010).

16. François, O. & Caye, K. Naturalgwas: An R package for evaluating genome-wide association methods with empirical data. Molecular Ecology Resources 18, 789–797 (2018).

17. Benjamini, Y. & Hochberg, Y. Controlling the False Discovery Rate: A Practical and Powerful Approach to Multiple Testing. Journal of the Royal Statistical Society Series B: Statistical Methodology 57, 289–300 (1995).

18. Alonso-Blanco, C. et al. 1,135 Genomes Reveal the Global Pattern of Polymorphism in Arabidopsis thaliana. Cell 166, 481–491 (2016).

19. Barber, R. F. & Candès, E. J. A knockoff filter for high-dimensional selective inference. Ann. Statist. 47, (2019).

20. He, Z. et al. GhostKnockoff inference empowers identification of putative causal variants in genome-wide association studies. Nat Commun 13, 7209 (2022).

